# Epidermal Growth Factor/c-Met receptor signalling crosstalk drives tunneling nanotube formation in A549 lung adenocarcinoma cells

**DOI:** 10.64898/2026.01.23.701283

**Authors:** Salonee Banerjee, Aya Elmeligy, Griselda Awanis, Natalia Cicovacki, Jack Scalcione, James McColl, Bertrand Lézé, Stefan Bidula, Jelena Gavrilovic, Derek Warren, Anastasia Sobolewski

## Abstract

Tunneling nanotubes (TNTs) are actin-based cytoplasmic connections that can mediate intercellular transfer of various cellular cargo and have been implicated in cancer progression and chemoresistance. However, the signalling mechanisms driving their formation remain poorly understood. Given the frequent dysregulation of EGFR and c-Met signalling in non-small cell lung cancer (NSCLC), and prior evidence of TNTs in lung adenocarcinoma patient samples, we investigated the role of EGFR and c-Met receptor signalling crosstalk in TNT induction in A549 lung adenocarcinoma cells. Stimulation with EGF, HGF, or in combination induced a concentration dependent increase in the formation of TNTs. TNTs exhibited typical characteristics, including F-actin expression, non-adherence to the substratum and facilitated intercellular trafficking of lysosomes, mitochondria, and lipid vesicles. EGFR was identified as a novel component of TNTs, but had little co-localisation with the c-Met receptor. Co-stimulation with HGF and EGF did not produce consistent additive or synergistic effects on TNT formation, suggesting shared downstream signalling. Furthermore, although EGFR and c-Met inhibition blocked EGF- and HGF-induced TNTs respectively, inhibition of both receptors was required to suppress TNTs following dual HGF/EGF treatment. Interestingly, blocking the EGF receptor alongside c-Met resulted in a more potent inhibition of HGF-induced TNTs, indicating crosstalk. Furthermore, inhibition of downstream MEK and PI3K pathways reduced HGF- or EGF- induced TNT formation, but dual inhibition was required to completely block TNT formation in HGF+EGF co-stimulated cells. These findings reveal a novel convergence of EGFR and c-Met and their downstream MAPK/PI3K pathways in TNT regulation, which can have important clinical implications in combinatorial receptor and cell signalling pathway targeting in NSCLC.

## INTRODUCTION

Non-small cell lung cancer (NSCLC) is the most common form of lung cancer, accounting for 85% of cases globally, with lung adenocarcinoma being the predominant subtype, accounting for 40% of all cases (Molina et al., 2008). Despite advancements in targeted therapies, NSCLC still has high mortality rates and poor prognosis due to its aggressive metastatic capabilities and frequent development of resistance to targeted therapies (Paez et al., 2004; Koulouris et al., 2022; Wilson et al, 2016; Liu et al., 2020).

Among the key oncogenic drivers of NSCLC are mutations in receptor tyrosine kinases (RTKs), notably including the epidermal growth factor receptor (EGFR; receptor for epidermal growth factor) and the c-Met receptor (receptor for hepatocyte growth factor) (Jorge, Kobayashi and Costa, 2014; Zhang et al., 2018). RTKs, upon ligand binding, undergo dimerisation or heterodimerisation, triggering autophosphorylation of intracellular tyrosine residues. This facilitates the recruitment of adaptor proteins and activation of downstream signalling pathways, which can promote cell proliferation, survival and motility (Du and Lovly, 2018; Uchikawa et al., 2021; Tito et al., 2025). EGFR mutations drive oncogenesis by constitutively activating these pathways (Yarden & Sliwkowski, 2001; Kosaka et al., 2014; Kim et al., 2014). Although EGFR tyrosine kinase inhibitors (TKIs) initially yield clinical responses, resistance frequently emerges (Abourehab et al., 2021). Third-generation EGFR-TKIs like osimertinib offer improved efficacy, yet resistance remains inevitable (Ramalingam et al., 2018; Cho et al., 2023). Similarly, dysregulation of HGF/c-Met signalling has been associated with cancer cell proliferation, epithelial-mesenchymal transition (EMT), invasion, and metastasis (Graziani et al., 1991; Ichimura et al., 1996; Corso et al., 2005; Sequist et al., 2011; Park et al., 2012; Maharati et al., 2022; Yao et al., 2024). Despite distinct ligands and receptors, EGFR and c-Met share downstream effectors such as the MAPK and PI3K pathways, and their crosstalk has been implicated in promoting aggressive tumour behaviour (Puri and Salgia, 2008; Acunzo et al., 2013; Fong et al., 2013; Crees et al., 2020). Moreover, resistance to EGFR-TKIs can occur through the activation of the c-Met pathway, often *via* c-Met amplification (Bean et al., 2007).

The complexity of the tumour microenvironment (TME) warrants better understanding of the mechanisms governing cell-to-cell communication, and the diverse factors that influence these processes to cause chemoresistance. Among these mechanisms, tunneling nanotubes (TNTs) have emerged as a novel and unconventional mode of intercellular connectivity. First described by Rustom et al. (2004) as thin, F-actin-based cytoplasmic protrusions capable of connecting cells over long distances, TNTs have since been identified in multiple cell types, reaching lengths of up to 350 µm (Ariazi et al., 2017; Cervantes and Zurzolo, 2021; Awanis et al., 2023). Their physiological relevance has also been demonstrated, with TNTs being observed in resected lung adenocarcinoma tissue obtained from patients (Lou et al., 2012). TNTs have since been increasingly implicated in tumour progression and chemoresistance, largely through the transfer of oncogenic cargo and subcellular organelles (Lou et al., 2012; Wang and Gerdes, 2015). In addition to organelle trafficking, TNTs have been shown to mediate the exchange of Ca²⁺ signals, extracellular materials, and oncogenes such as KRAS (Eugenin, Gaskill and Berman., 2009; Desir et al., 2019; Pepe et al., 2022),

Previous research on the mechanisms of TNT formation have predominantly focused on stress conditions (Arkwright et al., 2010; Wang et al., 2011; Lou et al., 2012), with limited understanding of whether frequently dysregulated factors and pathways in the TME are also involved in TNT formation. However, our recent work (Awanis et al., 2023) identified HGF/c-Met signalling as a promoter of TNT formation in A549 lung adenocarcinoma cells. Given the established roles of the EGF/EGFR and HGF/c-Met axes in NSCLC progression, in this study we address whether the EGFR/c-Met receptor crosstalk can contribute to TNT formation. This research provides further insights into the cell biology of how targeting these receptors in NSCLC may not only be working through effects on cancer cell growth (Ricciardi et al., 2010), but also through the formation of TNTs.

## RESULTS

### EGF, HGF, and EGF+HGF induce TNTs in A549 cells

In control conditions, A549 cells exhibited their characteristic cuboidal morphology, were adherent and grew in colonies (Fig. 1A). EGF, HGF and EGF+HGF treatment induced mesenchymal-like cell morphology. Notably, 100 ng/ml of EGF, HGF and EGF+HGF induced thin cytoplasmic protrusions (Fig. 1A) resembling TNTs described in the literature (Rustom et al., 2004) and earlier by us (Awanis et al., 2023). These TNTs were long and thin, either extending from one cell towards other cell(s) or forming a full connection between two or more distant cells.

**Fig 1.**
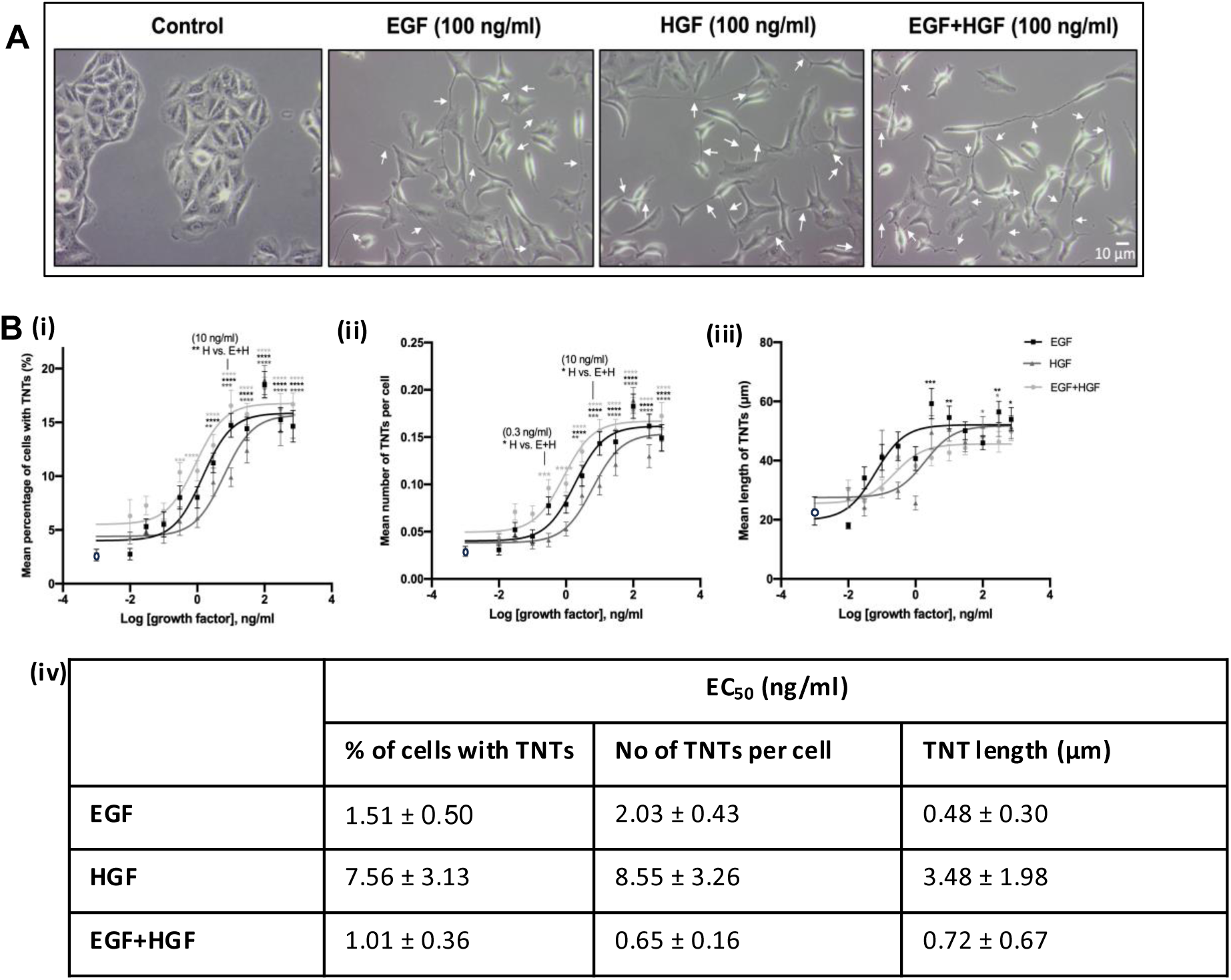
EGF, HGF and EGF+HGF induce tunneling nanotubes in a concentration dependent manner. **(A)** Representative phase contrast images demonstrate an increase in TNTs (white arrows) at the maximal concentration of EGF, HGF and EGF+HGF (100 ng/ml). The phase contrast images were quantified for TNT structures. Images were captured using 10x objective lens using an inverted microscope. **(B)** Log concentration-response curves for EGF (black), HGF (dark grey) and EGF+HGF (light grey) display the mean percentage **(i),** mean number of TNTs per cell **(ii),** and mean length of TNTs **(iii)** following 24 hours of growth factor treatment (0.01-700 ng/ml). Values are expressed as mean ± SEM, n = 3. ****P < 0.0001, ***P < 0.001, **P < 0.01 and *P < 0.05, compared to untreated control (denoted by white circle on graph). Statistical differences between growth factor treatments at specific concentrations are denoted by a vertical line, followed by the significance level and the treatments compared. **(iv)** EC_50_ values of EGF, HGF and EGF+HGF across the three TNT measurement parameters.

EGF, HGF and EGF+HGF-treated (0-700 ng/ml) A549 cells induced TNTs in a concentration-dependent manner (Fig. 1B). Quantitative analysis (mean ± SEM) showed only 2.68±0.53% of control cells had TNTs, with a mean of 0.03±0.005 TNTs per cell and a mean length of 22.97±4.81µm. Following EGF, HGF and EGF+HGF treatment, there was a concentration-dependent and significant increase in the mean percentage of cells with TNTs (Fig. 1Bi), mean number of TNTs per cell (Fig. 1Bii), and the mean length of TNTs (Fig. 1Biii). There was also a small leftward shift in the sigmoidal curves with EGF+HGF compared to individual EGF or HGF treatment, and with EGF compared to HGF, which is reflected in their respective EC_50_ values (Fig. 1Biv). There was also a significantly higher percentage of cells with TNTs with EGF+HGF compared to HGF treatment at 10 ng/ml (P < 0.01), and significantly higher number of TNTs per cell with EGF+HGF compared to HGF-treatment at 0.3 ng/ml and 10 ng/ml (P < 0.05). Besides 0.3 ng/ml and 10 ng/ml, the response with EGF+HGF treatment was comparable to individual treatment of EGF and HGF individually across the rest of the concentration range (n.s.). The percentage of cells exhibiting TNTs and the number of TNTs per cell reached their maximum threshold at 100 ng/ml for all three treatment conditions (EGF, HGF, and EGF+HGF). Beyond this concentration, the percentage, number of TNTs per cell and TNT length plateaued to similar levels with all three treatment conditions. While the mean lengths of 100 ng/ml EGF, HGF and EGF+HGF-induced TNTs were 46 µm, 52 µm and 48 µm respectively, the lengths spanned up to 270 µm, 257 µm and 244 µm long in EGF, HGF and EGF+HGF-treated cells respectively.

### HGF, EGF and HGF+EGF- induced TNTs express F-actin, are non-adherent and transfer cellular organelles

TNTs reported in the literature are shown to be comprised of F-actin, with some papers also reporting the presence of α-tubulin (Rustom et al., 2004; Huang et al., 2022), along with lack of adherence to the substrata and the cellular transport of organelles via TNTs (Rustom et al., 2004; Cervantes and Zurzolo, 2021; Awanis et al., 2023). EGF, HGF, and EGF+HGF stimulation induced TNTs that expressed F-actin throughout their lengths, with the thicker areas of TNTs expressing α-tubulin (Fig. 1Ai). F-actin expressing TNTs displayed non-adherence to the substrata, as shown in the 3-D reconstruction (Fig. 1Aii) and were maintained following trypsinization (Supplementary Fig. 1A). Scanning electron microcopy (SEM) showed a vast network of TNTs of different lengths following EGF, HGF and EGF+HGF treatment, with some TNTs extending over other cells to connect two or more distant cells (Fig. 2B). Occasionally, TNTs exhibited localised ‘bulges’ indicative of cellular cargo or vesicle-like structures where the TNT appeared to expand locally to accommodate the cargo (Fig. 2Biii).

**Figure 2.**
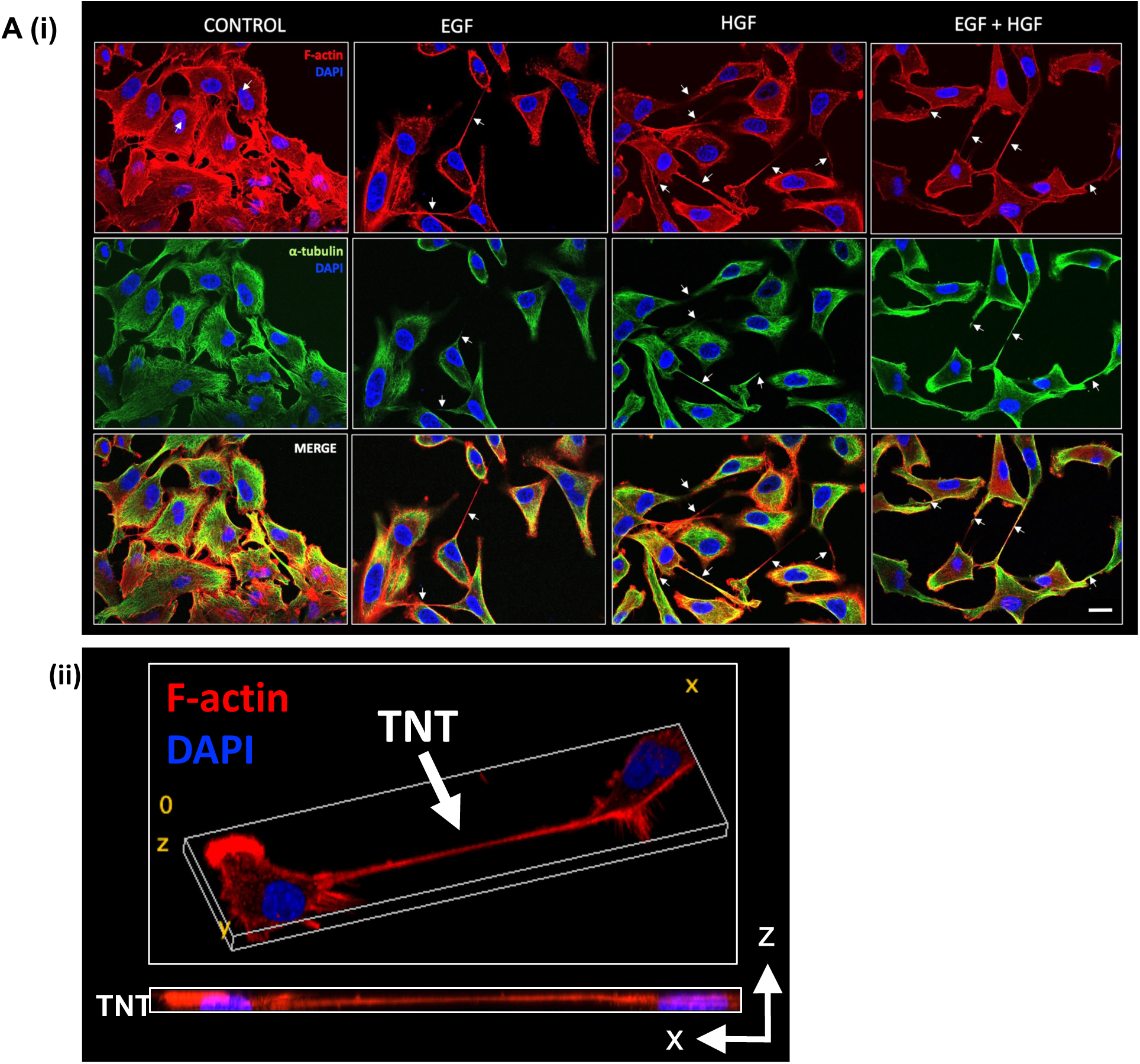
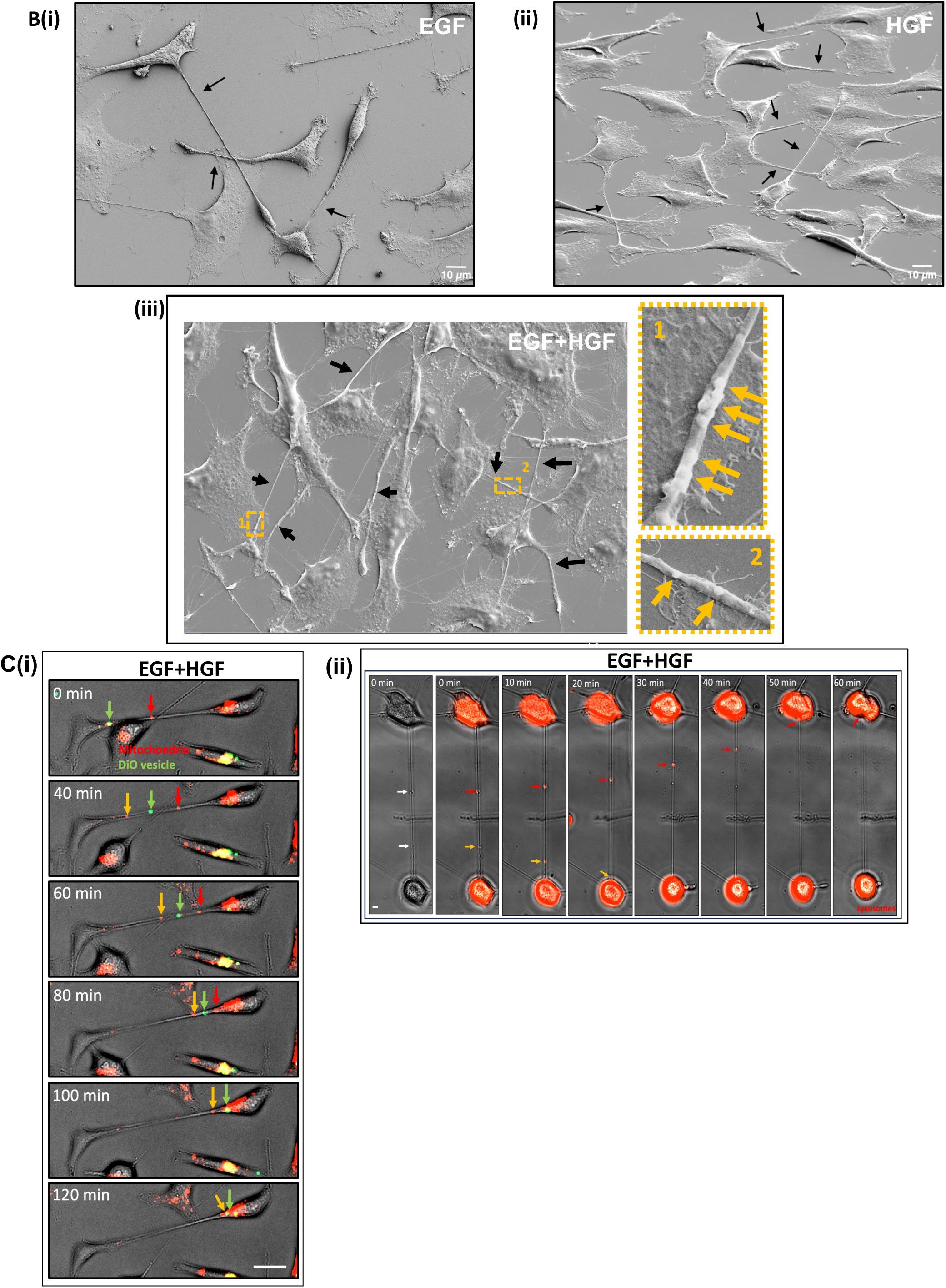
EGF, HGF and EGF+HGF-induced TNTs are non-adherent, express F-actin and α-tubulin, form vast networks and can transfer mitochondria, lipid vesicles and lysosomes (A)(i) Representative confocal images of control, EGF, HGF and EGF+HGF-treated cells (100 ng/ml) labelled with F-actin (red) and α-tubulin (green). White arrows denote TNTs. Confocal images were obtained at x40 magnification. Scale bar: 20 µm. (n=3) (ii) Three-dimensional reconstruction of an F-actin-positive non-adherent TNT induced by EGF+HGF. The orthogonal view demonstrates the xz projection of TNT hovering above the surface of the substratum. **(B)** Representative scanning electron micrographs demonstrate a vast network of TNTs in **(i)** EGF, **(ii)** HGF and **(iii)** EGF+HGF-treated A549 cells. The presence of cellular cargo within the EGF+HGF-induced TNTs is shown in the dotted zoomed panels (1) and (2) using yellow arrows. TNTs are denoted using black arrows. SEM images were obtained at magnifications ranging between x560 - x599. **(C)** Representative phase contrast and merged fluorescent images from a timelapse movie showing **(i)** two mitochondria (red and yellow arrows) and a DiO-labelled vesicle (green arrow) being trafficked within anEGF+HGF-induced TNT towards a red mitochondria-labelled cell. Scale bar: 10 µm and **(ii)** bidirectional transfer of lysosomes within an EGF+HGF-induced TNT (red and yellow arrows). Scale bar: 5 µm. Timelapse movies were obtained at 20x magnification. (n=3)

We have previously shown (Awanis et al., 2023) that HGF-induced TNTs can facilitate trafficking of mitochondria and lipid vesicles. We now show that EGF- (Supplementary fig. 1Bi and 1Bii) and HGF+EGF-induced TNTs (Fig. 2Ci and supplementary video 2Ci) can also traffic mitochondria and lipid vesicles. Mitochondria were trafficked unidirectionally (18.7 ± 4.87 %, 14.5 ± 4.37 % and 6.09 ± 2.24 % of TNTs induced by EGF, HGF, and EGF+HGF, respectively) and bidirectionally (1.14 ± 1.14 %, 3.78 ± 1.87 % and 6.09 ± 4.07 % of TNTs induced by EGF, HGF, and EGF+HGF respectively). Lipid vesicles were trafficked unidirectionally in 1.52 ± 1.52 %, 4.84 ± 2.03 % and 3.47 ± 2.45 % of TNTs induced by EGF, HGF, and EGF+HGF, respectively, and bidirectionally in 1.79 ± 1.79 % and 1.39 ± 1.39 % of HGF, and EGF+HGF-induced TNTs respectively (EGF N=129, HGF N=193, EGF+HGF N=161 for mitochondria/lipid vesicle transfer analysis). Figure 2Cii shows the bidirectional transfer of lysosomes between two EGF+HGF-treated cells over a period of 60 mins using time lapse microscopy (Supplementary video 2Cii). Similar transfer of lysosomes was observed in TNTs induced by EGF and HGF individually (Supplementary fig. 1Biii-iv). Further analysis showed that unidirectional lysosome transfer occurred in 12.3 ± 3.69 %, 13.8 ± 3.97 %, and 10.6 ± 3.98 % of TNTs induced by EGF, HGF, and EGF+HGF, respectively. Bidirectional lysosome transfer was less frequent, occurring in 1.33 ± 0.33 % of EGF-induced, 3.66 ± 2.06 % of HGF-induced and 1.25 ± 1.25% of EGF+HGF-induced TNTs (EGF N=106, HGF N=123, EGF+HGF N=85). No DiO or lysosomal transfer was detected in control TNTs, although unidirectional mitochondrial transfer occurred in 16% of control TNTs (Control N=33). Moreover, the mean rates of organelle transfer within TNTs were comparable across treatments for lysosomes (11.7, 10.7, and 8 nm/s), mitochondria (5.5, 9.2, and 12.7 nm/s), and DiO-labelled vesicles (2.2, 4.8, and 8.3 nm/s) for EGF, HGF, and EGF+HGF, respectively.

### C-Met and EGF receptor crosstalk regulates TNT formation in A549 cells

We next investigated the involvement of the EGFR and c-Met receptors in EGF/HGF/EGF+HGF-induced TNT formation using receptor inhibitors erlotinib and PHA-665752 respectively. Erlotinib inhibited the percentage and number of EGF-induced TNTs in a concentration-dependent manner, exhibiting a sigmoidal inhibition curve; with no effect on HGF and EGF+HGF-induced TNTs (Fig. 3Ai and ii). There was a gradual inhibition in the mean length of EGF-TNTs following erlotinib treatment, however these changes were not statistically significant (Fig. 3Aiii). The c-Met inhibitor PHA-665752 significantly reduced the percentage and number of HGF-induced TNTs in a concentration-dependent manner, exhibiting a sigmoidal inhibition curve (as we have previously shown (Awanis et al., 2023), but it had no effect on EGF and EGF+HGF-induced TNTs, (Fig. 3Bi and ii). There was a gradual reduction in the mean length of HGF-induced TNTs, although not statistically significant (Fig. 3Biii). In summary, while erlotinib selectively inhibited EGF-induced TNTs and c-Met inhibition selectively inhibited HGF-induced TNTs, the inhibition of either c-Met or EGFR did not affect EGF+HGF-induced TNTs.

**Figure 3.**
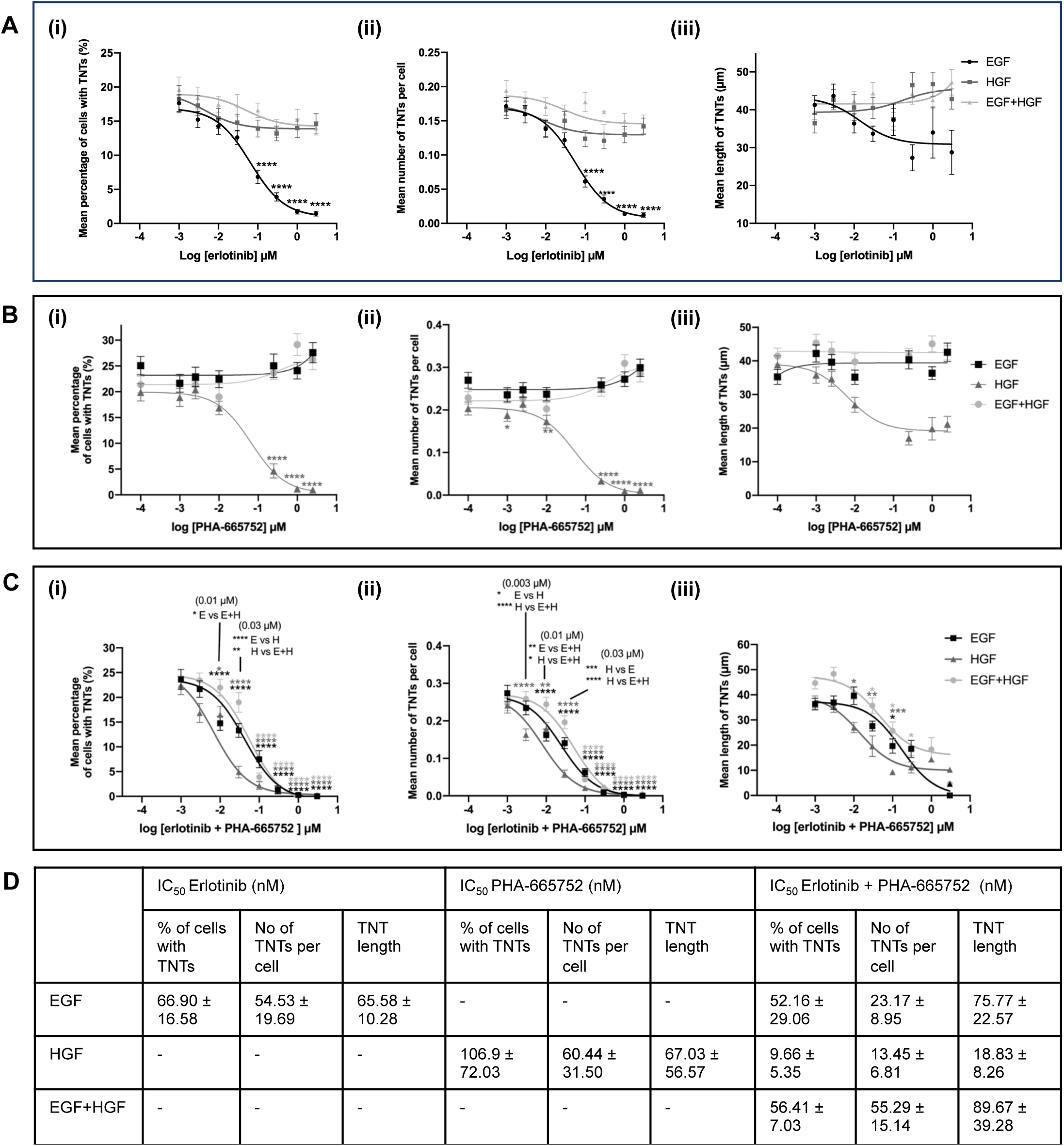
Effects of the C-met and EGF receptor inhibition on EGF, HGF and EGF+HGF-induced TNT formation in A549 cells. **(A)** EGF receptor (EGFR) inhibition using erlotinib (0.003 µM – 3 µM) inhibits EGF-induced TNTs in a concentration-dependent manner with no effect of EGFR inhibition on HGF and EGF+HGF-induced TNTs. The concentration-response curves demonstrate the (i) mean percentage of cells with TNTs (ii) mean number of TNTs per cell and (iii) mean TNT length. **(B)** c-Met inhibition using PHA-665752 (0.001 µM – 2.5 µM) inhibits HGF-induced TNTs in a concentration-dependent manner with no effect of c-Met inhibition on EGF and EGF+HGF-induced TNTs. The concentration-response curves demonstrate the (i) mean percentage of cells with TNTs (ii) mean number of TNTs per cell and (iii) mean TNT length. **(C)** Dual EGFR and c-Met receptor inhibition is required to abrogate EGF+HGF-induced TNTs. The concentration-response curves demonstrate the (i) mean percentage of cells with TNTs (ii) mean number of TNTs per cell and (iii) mean TNT length of cells treated with erlotinib+PHA-665752 (0.003 µM – 3 µM). **(D)** IC_50_ values of erlotinib, PHA-665652 and erlotinib+PHA-665752 in inhibiting TNT parameters of EGF, HGF and EGF+HGF-induced TNTs. Values are expressed as mean ± SEM, n = 3. ****P < 0.0001, ***P < 0.001, **P < 0.01 and *P < 0.05, compared to untreated (no inhibitor) controls for EGF (black), HGF (dark grey), or EGF + HGF (light grey). Statistical differences between growth factor treatments at specific concentrations of erlotinib + PHA-665752 are denoted by a vertical line, followed by the significance level and the treatments compared.

Simultaneous blocking of the EGF and c-Met receptor (Fig. 3Ci-iii) was required to inhibit EGF+HGF-induced TNTs and this occurred in a concentration-dependent manner with a significant reduction in the mean percentage, the mean number and the mean length. EGF or HGF-induced TNTs also showed sigmoidal concentration-dependent inhibition curves with EGFR/c-Met dual blocking. Interestingly, dual EGFR/c-Met inhibition was more effective in inhibiting HGF-induced TNTs than c-Met inhibition alone, as evidenced by the leftward shift in the HGF curve and the corresponding IC_50_ values (106.9 nM with PHA-665752 alone vs 9.66 nM with PHA-665752+erlotinib). Similarly, but not as marked, dual EGFR/c-Met inhibition was more effective in inhibiting EGF-induced TNTs than EGFR inhibition alone (66.9 nM with erlotinib alone vs 52.16 nM with erlotinib+PHA-665752) (Fig. 3D).

### TNTs show distinct localisation of the EGFR and c-Met receptor

Immunofluorescence labelling of EGFR and c-Met showed that EGFR (green) and c-Met (red) were co-localised at the cell membrane in the epithelial colonies in control cells (Fig. 4A). HGF, EGF+HGF treatment, but not EGF induced the internalisation of c-Met (red) from the cell membrane. C-Met was ubiquitously expressed along the length of EGF, HGF and HGF+EGF induced TNTs. We previously showed that C-Met is expressed in HGF-induced TNTs (Awanis et al., 2023). EGF and EGF+HGF treatment caused the internalisation of EGFR (green) into distinct puncta that were expressed in the cell. Unlike c-Met, not all TNTs expressed EGFR, with puncta present in 60%, 53%, 83% and 35% of control, EGF, HGF and EGF+HGF-induced TNTs respectively (Fig. 4A).

**Figure 4.**
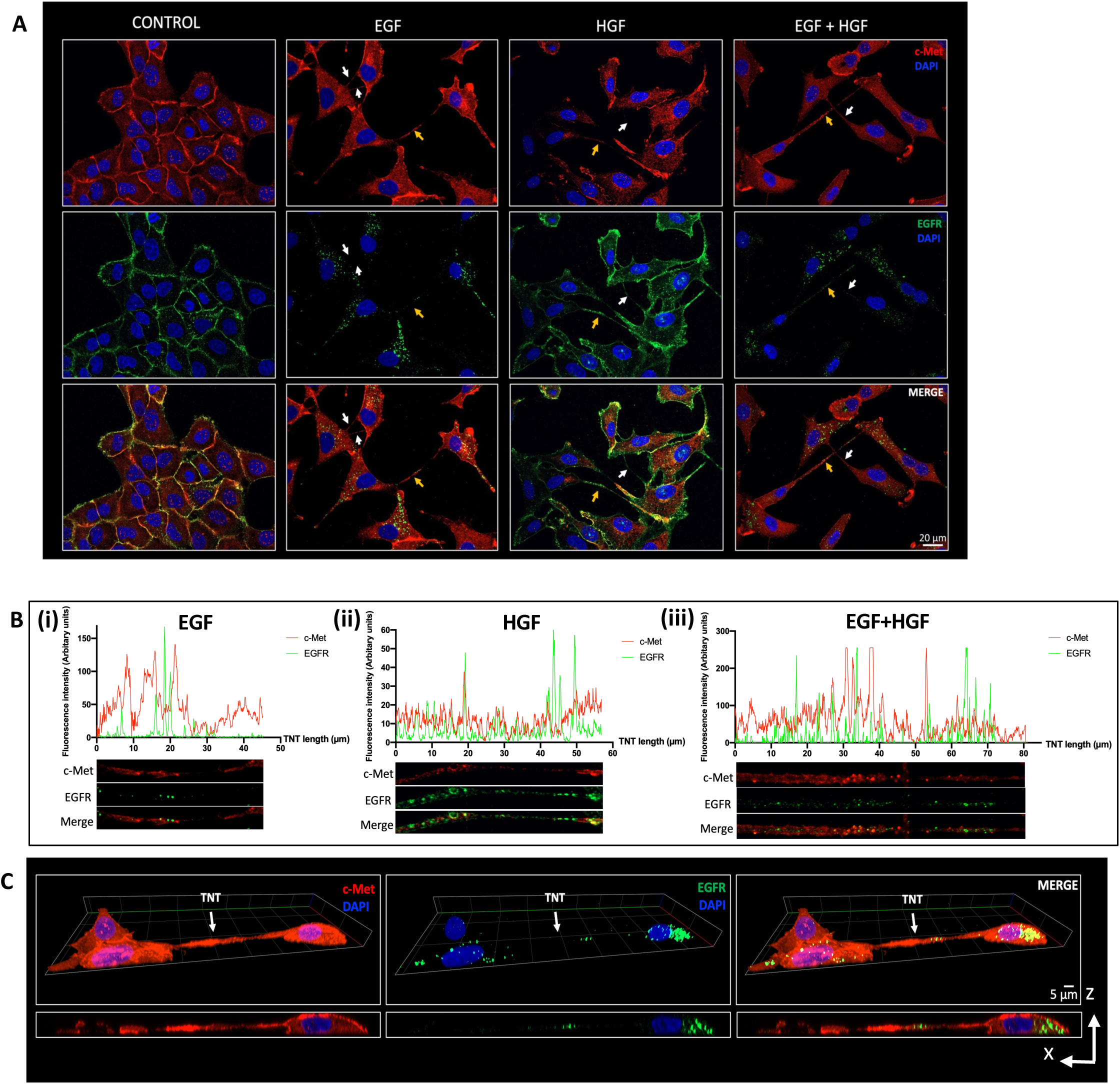
EGF, HGF and EGF+HGF-induced TNTs express the EGF and c-Met receptors. **(A)** Representative confocal images showing expression of c-Met (red) and EGFR (green) along the observed TNTs (white arrows). TNTs chosen for line scan are indicated using yellow arrows **(B)** Representative line scans demonstrating the expression of c-Met and EGFR along **(i)** EGF **(ii)** HGF and **(iii)** EGF+HGF-induced TNTs. **(C)** Representative 3D reconstruction images showing an example of a non-adherent EGF+HGF-induced TNT expressing c-Met (red) and EGFR (green). The Z-stack was obtained using ZEN lite software with optimal Z-step of 0.21 µm. Confocal images were obtained at x40 magnification. (n=3).

There was minimal spatial overlap of EGFR and c-Met puncta, which was reflected in the line-scan profiles showing the distribution of EGFR and c-Met along the length of the TNTs (Fig. 4B). Co-localisation analysis of EGFR and c-Met revealed a Pearson’s correlation coefficient (PCC) value of 0.219 for control, 0.132 for EGF, 0.221 for HGF and 0.244 for EGF+HGF-induced TNTs (Control N=10, EGF N=16, HGF N=12, EGF+HGF=25). This confirmed that there was negligible weak co-localisation between c-Met and EGFR in the TNTs. An xz 3D reconstruction and orthogonal view using confocal z stack images demonstrates the non-adherence of EGFR/c-Met expressing TNTs (Fig. 4C), and similar non-adherence was observed across all conditions as we observed earlier. Localisation of the EGF/c-Met receptors within lysosome or early endosome subcellular compartments in TNTs was next investigated by co-labelling for Lamp-1 and EEA1. Lamp-1 was present in all TNTs (Supplementary fig. 2Ai) but did not co-localise with c-Met and EGFR (Supplementary Fig. 2Aii-iv). EEA1 was also found in TNTs (Supplementary Fig. 2B) with occasional co-localisation with c-Met or EGFR in EGF, HGF and EGF+HGF-induced TNTs (Supplementary Fig. 2Bii-iv).

### Convergence of EGFR and c-Met receptor signalling at the MAPK and PI3K pathways drives TNT formation

To evaluate the roles of the MEK/MAPK and PI3K pathways in TNT formation, cells were treated with MEK inhibitor PD98059, PI3K inhibitor LY294002, or both, at various concentrations (0.3-100 µM) in the presence of HGF, EGF, or EGF+HGF. Both inhibitors reduced TNT formation in a concentration-dependent manner across all growth factor conditions (Fig. 5). MEK inhibition using PD98059 significantly inhibited TNT formation, affecting both the percentage of cells with TNTs and number of TNTs per cell, with HGF-induced TNTs showing the greater sensitivity to the drug. However, EGF+HGF-treated cells required a higher PD98059 concentration (30 µM) for effective TNT inhibition (Fig. 5Ai and ii). PD98059 significantly reduced the length of EGF+HGF-induced TNTs at 100 µM, with no effect observed in the length of EGF or HGF-induced TNTs (5Aiii). LY294002 also reduced TNT formation in a dose-dependent manner evidenced by the inhibition of all three parameters, showing a comparable effect to PD98059 in inhibiting EGF and HGF-induced TNTs but less effective inhibition of EGF+HGF-induced TNTs. Higher concentrations of LY294002 (30 µM) were required to effectively inhibit EGF + HGF-induced TNT formation compared to the concentrations needed for TNT inhibition with individual EGF or HGF stimulation, a pattern previously observed with PD98059 treatment alone (Fig. 5Bi-iii). LY294002 significantly reduced the length of EGF and HGF-induced TNTs at 10-30 µM, while 100 µM was required to inhibit EGF+HGF-induced TNT length (Fig. 5Biii). Dual treatment with PD98059 and LY294002 produced a stronger inhibitory effect on TNT formation, with pronounced inhibition of percentage of cells with TNTs and number of TNTs per cell at lower doses compared to singular drug treatment, especially in EGF+HGF conditions (Fig. 5Ci and ii). The respective IC_50_ values confirmed that combined MEK/PI3K inhibition was most effective in TNT abrogation, with the lowest IC_50_ observed with dual inhibitor treatment (3.13 µM) compared to only MEK (13.12 µM) or PI3K (17.45 µM) inhibition of EGF+HGF-induced TNTs (Fig. 5D).

**Fig 5.**
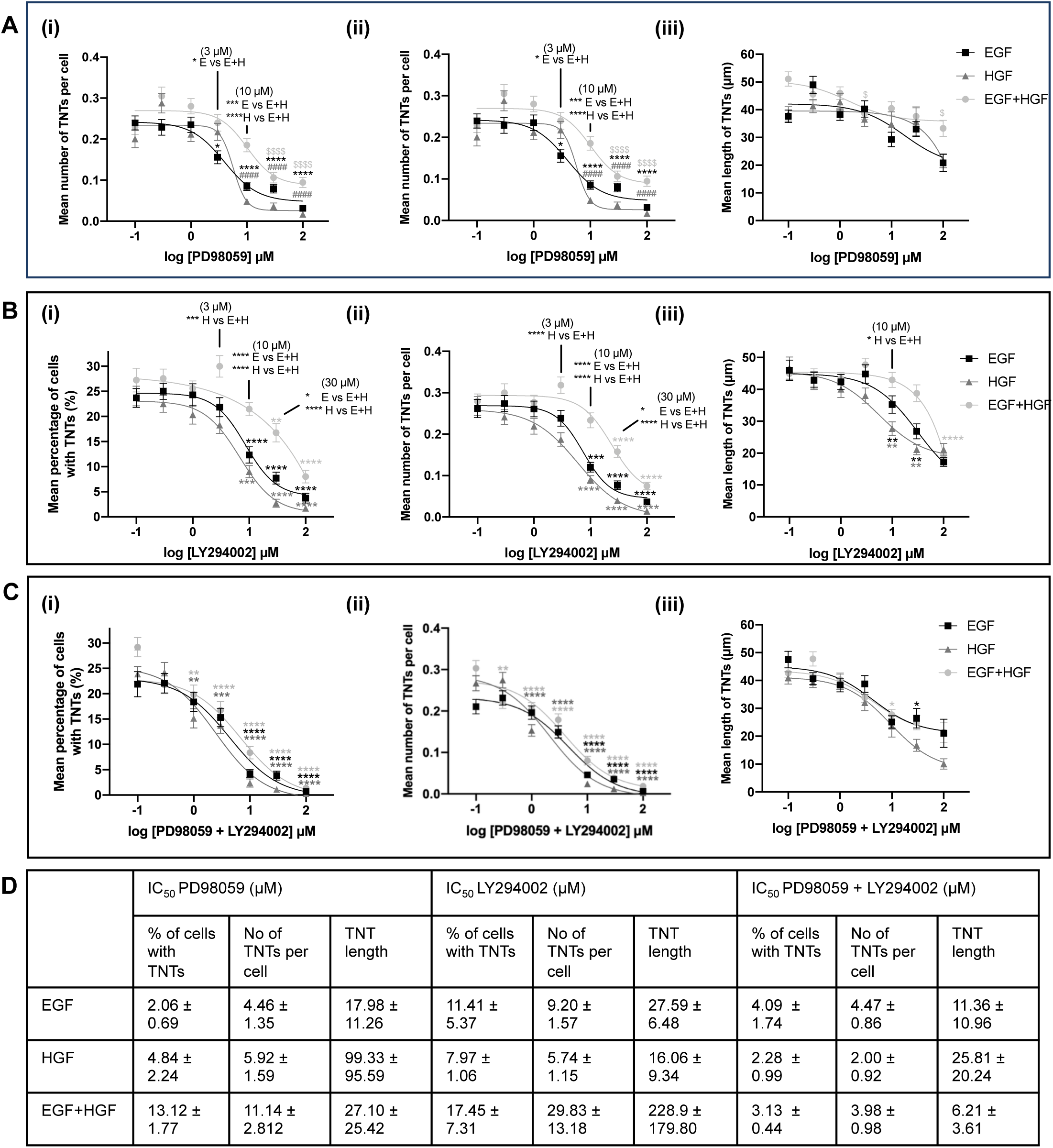
The MEK/MAPK and PI3K pathways mediate HGF, EGF and EGF+HGF-induced TNT formation in A549 cells. (A) MEK inhibition using PD98059 (0.1 µM – 100 µM) inhibits TNTs induced by EGF (black), HGF (dark grey) and EGF+HGF (light grey) (100 ng/ml). The dose-response curves demonstrate the decrease in (i) mean percentage of cells with TNTs and (ii) mean number of TNTs per cell, with no significant decrease in (iii) mean TNT length across the concentration range. (B) PI3K inhibition using LY294002 (0.1 µM – 100 µM) inhibits TNTs induced by EGF, HGF and EGF+HGF (100 ng/ml). The dose-response curves demonstrate the decrease in (i) mean percentage of cells with TNTs (ii) mean number of TNTs per cell, and (iii) mean TNT length. (C) Dual MEK and PI3K inhibition using PD98059+LY294002 (0.1 µM – 100 µM) suppressed TNTs induced by EGF, HGF and EGF+HGF (100 ng/ml) in a concentration-dependent manner. The dose-response curves demonstrate the decrease in (i) mean percentage of cells with TNTs (ii) mean number of TNTs per cell, with no significant difference in (iii) mean TNT length. **(D)** IC_50_ values of PD98059, LY294002 and PD98059+LY294002. Values are expressed as mean ± SEM, n = 3. Statistical significance (****P < 0.0001, ***P < 0.001, **P < 0.01, *P < 0.05) indicates comparisons to the respective untreated controls for each growth factor condition: EGF (black), HGF (dark grey), and EGF+HGF (light grey)- i.e., inhibitor-treated samples were compared to the corresponding no-inhibitor controls containing the same growth factor. Statistical differences between growth factor treatments at specific concentrations are denoted by a vertical line, followed by the significance level and the treatments compared.

## DISCUSSION

By recapitulating *in vitro* the combined presence of EGF and HGF within the lung tumour microenvironment, this study identifies a novel role for EGF and c-Met receptor signalling crosstalk in the induction of TNTs in A549 cells. Stimulation with EGF, HGF, or EGF+HGF promoted the formation of TNTs that were F-actin-rich, non-adherent, and capable of transporting lysosomes, mitochondria and DiO-labelled lipid vesicles. Moreover, c-Met, EGFR, EEA1 and Lamp-1 were identified as components of TNTs. Inhibition of the c-Met receptor or EGFR selectively inhibited HGF or EGF-induced TNTs respectively, but dual inhibition was required to inhibit EGF+HGF-induced TNTs. Moreover, the dual inhibition of both the downstream PI3K and MAPK pathways was required to strongly inhibit EGF+HGF-induced TNTs, while single PI3K or MAPK inhibition was sufficient for the maximal abrogation of EGF- or HGF-induced TNTs.

The structural and functional characteristics of the TNTs induced by EGF, HGF, and EGF+HGF were consistent with canonical characteristics of TNTs reported in the literature; i.e. F-actin expression, non-adherence, and ability for long distance intercellular transport of cellular materials (Rustom et al., 2004; Lou et al., 2012; Pasquier et al., 2012; Osswald et al., 2015; Cole et al., 2021). Consistent with our findings, the expression of F-actin and α-tubulin in TNTs has been reported in several cell types, with α-tubulin being particularly abundant in thicker or more stable TNTs (Gerdes and Carvalho, 2008; Melwani and Pandey, 2023). Moreover, scanning electron microscopy revealed membrane bulges along TNTs consistent with previously described ‘gondolas’, which have been suggested to facilitate lipid and protein transport in other cells (Hurtig et al., 2010; Franco et al., 2020). While lysosomal trafficking *via* TNTs has been documented in other cells (Rustom et al., 2004; Önfelt et al., 2006; Dilsizoglu Senol et al., 2021; Omsland et al., 2017), we now report the uni- and bi-directional transfer of lysosomes *via* TNTs in A549 cells. This is particularly important given the role of lysosomes in autophagy and drug sequestration in cancer (Kallunki, Olsen and Jäättelä, 2013; Hraběta et al., 2020). TNTs also exhibited the transfer of mitochondria and lipid vesicles. Mitochondrial transfer *via* TNTs has been previously shown in several cell lines (Onfelt et al., 2006; Wang and Gerdes, 2015; Jackson et al., 2016; Yang et al., 2022; Chakraborty et al., 2023) and in lung adenocarcinoma tumours (Lou et al., 2012), and has implications in rescue and chemoresistance mechanisms between cancer cells (Warburg, 1956; Spees et al., 2006; Wang and Gerdes, 2015; Yang et al., 2022). An emerging question is whether organelle trafficking via TNTs is selective and influenced by TNT subtype or environmental stressors. TNTs are structurally heterogeneous, with thicker, microtubule-containing TNTs supporting transport of large organelles (Zamberlan and Semenzato, 2025). Increasing evidence also indicates preferential organelle transfer under cellular stress. For example, FAK-deficient squamous cell carcinoma cells transfer lysosomes to FAK-proficient cells via TNTs, facilitating adaptation to impaired signalling (Sáenz-de-Santa-María et al., 2017). Similarly, TNT-mediated mitochondrial transfer has been shown to support cell survival and metabolic reprogramming by enabling the transfer of functional mitochondria from healthy donor cells to recipient cells with compromised mitochondrial function or undergoing apoptosis (Spees et al., 2006; Wang and Gerdes, 2015)

A novel observation was also the distinct localisation of EGFR and c-Met within TNTs. Despite their co-localisation at the plasma membrane, these receptors did not strongly co-localise within TNTs under any condition. Co-localisation between the EGFR and c-Met receptor has been documented widely in cancer including in NSCLC cells and patient biopsies (Velpula et al., 2012; Ortiz-Zapater et al., 2017; Crees et al., 2020; Harwardt et al., 2020), however their co-expression in TNTs was unexplored prior to this study. We also observed occasional co-localisation of EGFR and c-Met with EEA1 within TNTs, aligning with EEA1’s established role in early endocytic trafficking of these receptors (Leonard et al., 2008; Sorkin & Goh., 2009; Barrow-McGee & Kermorgant., 2014; Viticchiè & Muller., 2015). This observation raises intriguing questions regarding how growth factor receptors are trafficked through TNTs and represents an important direction for future investigation.

TNT formation by EGF or HGF stimulation occurred in a concentration-dependent manner. The effective concentrations required for TNT induction were similar to those that have been shown to promote cell proliferation (Puri and Salgia., 2008), migration (Lu et al., 2001; Kermorgant et al., 2004; Harrison et al., 2013; Schelch et al., 2021), and invasion (Lu et al., 2001; Harrison et al., 2013). HGF induced TNT formation with a maximal effect observed at 100 ng/ml, consistent with our previous findings (Awanis et al., 2023; Chen and Zhao, 2024). We now show that EGF is also a potent inducer of TNTs in A549 cells, with maximal response at 100 ng/ml. Previous studies have shown EGF to induce TNTs in various cell types (Lou et al., 2012; Carter et al., 2019; Hanna et al., 2019; Cole, Dahl and Dahl, 2021), at comparable EGF concentration ranges to our experiments EGF. A leftward shift in the curve with EGF+HGF compared to EGF or HGF stimulation suggested a marginal increase in potency. The subtle shift may reflect partial convergence or amplification of signalling pathways, possibly through receptor crosstalk or transactivation, as previously reported in NSCLC (Puri and Salgia., 2008; Tang et al., 2008). However, there was no consistent difference of EGF+HGF across the full range of concentrations compared to single EGF or HGF treatment. In contrast to our findings in TNTs, EGF+HGF co-stimulation has been shown to synergistically enhance proliferation in NSCLC and colorectal cancer cells (Puri and Salgia, 2008; Liska et al., 2011) and invasion in ovarian cancer models (Zhou et al., 2007), suggesting that the cell signalling pathways governing TNT formation are regulated differently from those driving proliferation or invasion.

The role of c-Met receptor signalling has been demonstrated in HGF-induced TNT formation (Awanis et al., 2023), and our findings now show that EGFR signalling also drives TNT induction in A549 cells. Moreover, the dual inhibition of EGFR and c-Met was not only required to suppress EGF+HGF-induced TNTs, but also increased the sensitivity of cells to EGF- or HGF-induced TNT inhibition, compared to single-receptor targeting. This suggested a signalling crosstalk between the EGFR and c-Met receptors, which extends from previous findings of EGFR/c-Met interplay in promoting proliferation and tumour growth, particularly in NSCLC (Blazek, Bonomi and Plate, 2008; Puri and Salgia, 2008; Lal et al., 2009). Notably, c-Met activation is also a well-characterised bypass mechanism in EGFR-TKI-resistant tumours, allowing sustained downstream signalling despite EGFR blockade (Engelman et al., 2007; Yu et al., 2013; Papadimitrakopoulou et al., 2018). It is worth noting that the doses of the EGFR and/or c-Met inhibitor required to suppress TNT formation in our study were markedly lower (in the nM range) than those typically needed to inhibit other oncogenic behaviours such as proliferation or migration (µM) (Kritikou et al., 2013; Glaser et al., 2021; Alhazzani et al., 2023; Matsubara et al., 2010; Yang et al., 2008). This suggests that TNT formation may be more sensitive to the inhibition of EGFR/c-Met signalling than other cellular responses, indicating that TNTs could be an early and highly responsive adaptation mechanism to TME stimuli. As such, lower doses of dual c-Met/EGFR inhibitor treatment may be sufficient to disrupt tumour cell communication *via* TNTs while minimising cytotoxic effects to healthy cells. Interestingly, an EGFR-Met-targeting biospecific antibody amivantamab, has recently shown promise in clinical trials (Cho et al., 2023), followed by its approval for treatment of EGFR-mutant NSCLC. It may be possible that TNTs have been overlooked as a possible mechanism of bypass signalling, which can be averted by EGFR-c-Met dual inhibition.

MEK/MAPK and PI3K/AKT are well-known signalling cascades acting downstream of EGFR/c-Met to regulate proliferation, survival, and motility in cancer cells (Mendoza, Er and Blenis, 2011). We now show that EGF/HGF-induced TNT formation is also driven by these pathways. MEK inhibition was also more effective than PI3K inhibition at suppressing EGF+HGF-induced TNTs, resembling previous observations in NRAS-mutant melanoma cells, where MEK inhibition caused greater growth suppression than PI3K, AKT, or mTOR inhibition (Posch et al., 2013). Furthermore, only combined MEK+PI3K inhibition fully abrogated EGF+HGF-induced TNTs, indicating that EGFR and c-Met converge on these pathways to drive TNT formation. This also suggests that dual receptor activation produces additive activation of both MAPK and PI3K signalling, as reflected by the markedly lower IC₅₀ values observed with dual inhibition compared to inhibition of either pathway alone. There has been previous evidence of compensatory MEK/ERK activation following PI3K/AKT/mTOR blockade (Carracedo et al., 2008; Serra et al., 2021) and also the ability of combined MEK/MAPK and PI3K/AKT targeting to overcome EGFR-TKI resistance in NSCLC (Li et al., 2011). Moreover, MAPK/PI3K co-inhibition has been shown to synergistically suppress tumour proliferation in colorectal cancer (Haagensen et al., 2012), skin cancer (Temblador et al., 2022), and NSCLC (Li et al., 2011; Heavey et al., 2016; Qu et al., 2021; Liu et al., 2024). Our findings extend this observation in the context of TNTs, highlighting a downstream convergence of EGFR and c-Met signalling via the activation of both the MAPK and PI3K signalling cascades, where the simultaneous inhibition of both pathways is required to disrupt this adaptive TNT network.

Understanding the signalling mechanisms driving TNT formation can be crucial for understanding tumour progression, particularly in NSCLC where resistance to targeted therapies remains a major challenge. Although EGFR and c-Met TKIs, and more recently, bispecific dual inhibitors, are in clinical use, their downstream mechanisms in modulating TNT-mediated communication had not been previously explored. Our findings have revealed that EGF/EGFR and HGF/c-Met receptor crosstalk extends beyond classical oncogenic proliferative pathways and has a crucial role in TNT formation, suggesting their broader role in shaping cancer cell behaviour and most importantly chemoresistance. This research could potentially inform NSCLC cancer treatment regimens: Following conventional drug therapies, lower dose combinatorial EGFR/c-Met inhibitor treatments could then be used to inhibit TNT formation and prevent chemoresistance.

## MATERIALS AND METHODS

### Reagents

#### Cytokines and Inhibitors

HGF and EGF were both purchased from PeproTech. The EGFR inhibitor (Erlotinib, 0.003 µM - 3 µM), c-Met inhibitor (PHA-665752, 2.5nM - 1µM) and MEK pathway inhibitor (PD98059, 0.3 µM - 100 µM) were obtained from Cayman Chemicals. The PI3K pathway inhibitor (LY294002, 0.3 µM - 100 µM) was acquired from Selleck Chemicals. DMSO was acquired from Sigma.

#### Immunofluorescence

Rabbit anti-Met, rat anti-EGFR and rat anti-α-tubulin were obtained from Abcam. Mouse anti-EEA1 and mouse anti-LAMP1 were purchased from ProteinTech. LysoTracker™ Red DND-99, Rhodamine Phalloidin, DiO’ and DiOC18(3) (3,3’-Dioctadecyloxacarbocyanine Perchlorate) and the Alexa-Flour 488, 568 and 647-conjugated secondary antibodies were all obtained from Thermofisher Scientific. The mitochondrial staining kit, red fluorescence-cytopainter, was purchased from Abcam. Vectashield mounting medium was obtained from Vector Laboratories Ltd. DAPI was obtained from Invitrogen.

#### Cell Culture

The A549 human lung adenocarcinoma cell line was obtained from the America Type Culture Collection (ATCC) and grown in Dulbecco’s modified Eagle’s medium (DMEM) media (Gibco) supplemented with 10% fetal bovine serum (FBS), 20 mM L-glutamine, 10,000 U/ml penicillin, and 10,000 μg/ml streptomycin (Gibco). Cells were maintained in a humidified atmosphere at 37°C at 5% CO_2_ levels. For growth factor treatment of A549 cells, DMEM media supplemented with 2% FBS, 20 mM L-glutamine, 10,000 U/ml penicillin, and 10,000 μg/ml streptomycin. A549 cells were seeded at 5x10^3^ cells per well using 10% FBS supplemented DMEM Media in a 12 well plate and incubated for 72 hours in a humidified atmosphere. After incubation, cells were treated with HGF (0.01–700 ng/ml) and/or EGF (0.01–700 ng/ml) in 2% FBS supplemented DMEM media and incubated for a further 24 hours before phase contrast imaging. For inhibitor studies, A549 cells were treated with inhibitors detailed above individually or in combination (in the case of PHA-665752+erlotinib; PD98059+LY294002) in 2% FBS supplemented DMEM media or their corresponding vehicle control in 2% FBS supplemented DMEM media for 30 minutes before being treated with 100ng/ml of HGF and/or 100ng/ml of EGF. Cells were subsequently incubated for 24 hours before phase contrast imaging. The concentration range of inhibitors used did not affect cell viability when an AlamarBlue assay was undertaken.

#### TNT retainment trypsinization experiments

To demonstrate the TNTs are retained post-trypsinization, A549 cells were seeded at 5x10^3^ cells per well in the 10% supplemented DMEM media and incubated for 72 hours. They were then treated with 100ng/ml of HGF and/or 100ng/ml of EGF in 2% supplemented FBS media and incubated for a further 24 hours. Cells were then directly treated with 500µl of trypsin (Gibco) and recorded using GXCAM3EY-5 camera at x10 magnification on a Zeiss Primo Vert Inverted White Light Microscope using GXCapture-T software until cells were fully detached.

#### Immunofluorescence labelling and confocal microscopy

A549 cells were seeded at 7.5x10^3^ cells per well in a 12 well plate containing 19 mm No.0 glass coverslips (Thermo Fisher Scientific) and incubated for 72 hours before treatment with 100ng/ml of HGF and/or 100ng/ml of EGF for a further 24 hours. For c-Met and EGFR immunolabelling, cells were fixed with 100% ice-cold methanol for 10 minutes, permeabilised with 0.2% Triton-X for 10 minutes and blocked with 3% BSA in PBS for 45 minutes. Cells were then incubated with primary antibodies to c-Met and EGFR (1:100 and 1:50 respectively) in 3% BSA overnight at 4°C. For α-tubulin immunolabelling, cells were fixed with 4% PFA for 10 minutes, permeabilised with 0.5% Triton-X for 3 ½ minutes and then blocked with 3% BSA in PBS for 45 minutes. Cells were then incubated overnight at 4°C with the anti-α-tubulin primary antibody (1:100) in 3% BSA. For EEA1 and LAMP1 co-labelling with EGFR and c-Met, cells were fixed with 100% ice-cold methanol for 10 minutes, permeabilised with 0.2% Triton-X for 10 minutes and blocked with 5% goat serum for 1 hour. Cells were then incubated with EEA1/LAMP1 (1:100), EGFR (1:50) and c-Met (1:100) primary antibodies in 3% BSA overnight at 4°C. Immunolabelled proteins were then incubated with respective species-specific Alexafluor-conjugated secondary antibodies (488, 568, and 647 nm) for 2 hours at 4°C in the dark. Cells were counterstained with DAPI and then imaged with the LSM980-Airyscan confocal microscope (Carl Zeiss, Germany) using 40x 1.3 NA oil objective with a pinhole setting of 1 AU.

#### Scanning Electron Microscopy

A549 cells were seeded at 7.5x10^3^ cells per well in a 12 well plate containing 12mm No.1 coverslips (VWR) that were coated with Poly-L-Lysine (0.01% w/v; Sigma-Aldrich) and incubated for 72 hours before treatment with 100ng/ml of HGF and/or 100ng/ml of EGF for a further 24 hours. Cells were fixed with 2.5% glutaraldehyde in 0.1M cacodylate buffer (both Sigma-Aldrich) at 4°C overnight before being washed twice with 0.1M cacodylate buffer and post-fixed with 2% osmium tetroxide (Sigma-Aldrich) for an hour in the dark at 4°C. Cells were subsequently washed with 0.1 M cacodylate buffer and dehydrated gradually through an ethanol series (25%, 50%, 75% and 100%; 10 minutes per step) and left to air dry overnight. Prepared samples on coverslips were mounted on specimen stubs and sputter coated with gold before being imaged using a Gemini SEM300 microscope (Carl Zeiss, Germany) at a 60° angle and viewed at 10 kV.

#### Live Cell trafficking of organelles

For the visualisation of lysosome transfer along TNTs, A549 cells were seeded at 7.5x10^3^ cells per well on glass chamber slides (Ibidi) and incubated for 72 hours before being treated with 100ng/ml of HGF and/or 100ng/ml of EGF and a further incubation for 24 hours. Cells were then treated with lysotracker (5μM) for 30 minutes before imaging on a Zeiss Axio Observer 7 microscope using a 20x 0.5 NA objective every 10 minutes for at least 10 hours. For the visualisation of mitochondria and lipid vesicles, two separate cell populations were treated with DiO (5µM) or red mitochondrial cytopainter (according to manufacturer’s instructions) for 30 minutes before being seeded together at a 1:1 ratio on glass chamber slides. The co-seeded cells were then treated with HGF and/or EGF for a further 24 hours. Live cells were then imaged using a Zeiss Axio Observer 7 microscope using a 20x 0.5 NA objective every 10 minutes for at least 10 hours.

### TNT analysis

#### Phase contrast image analysis

Phase contrast images were captured on a Zeiss Primo Vert Inverted White Light Microscope using GXCapture-T software and GXCAM3EY-5 camera. A 10x objective was used to capture 10 images per well. The ImageJ software was used to count the number of cells with TNTs to calculate the percentage of cells with TNTs. Additionally, the number of TNTs per cell and the length (μm) of all TNTs was measured from the narrow protrusion point of the TNT-initiating cells to the end of the TNTs. **Confocal image analysis:** Non-adherence of F-actin-based TNTs and XZ-localisation of EGFR and c-Met in TNTs was assessed by acquiring Z-stacks at 0.21–0.27 μm intervals. The ZEN lite and ImageJ software was used to obtain the 3D reconstruction and orthogonal views. The co-localisation analysis between EGFR and c-Met, in TNTs was performed using the JACoP (Just Another Co-localization Plugin) in the Fiji ImageJ software (Bolte and Cordelières, 2006) using Z-stack images (with Z steps between 0.21-0.55 µm). Each Z-plane was considered separately in the calculation of the Pearson’s correlation coefficient (PCC) to determine co-localisation. Line scans were plotted by generating fluorescence intensity profiles of c-Met and EGFR along the length of the TNTs using ImageJ.

#### Statistical analysis

Experiments were independently performed at least 3 times, with at least 600 cells analysed per condition in total. All data was expressed as mean ± SEM for histograms and log dose-response curves comparing control to growth factor-treated/inhibitor-treated groups. One-way ANOVA and Tukey’s multiple comparison tests on GraphPad Prism 8 were used to determine statistical significance. P < 0.05 was set as the cut-off value for statistical significance.

