## Supplementary figures 1 and 2 for "Epidermal Growth Factor/c-Met receptor signalling crosstalk drives tunneling nanotube formation in A549 lung adenocarcinoma cells"

A(i)

(ii)

(iii)

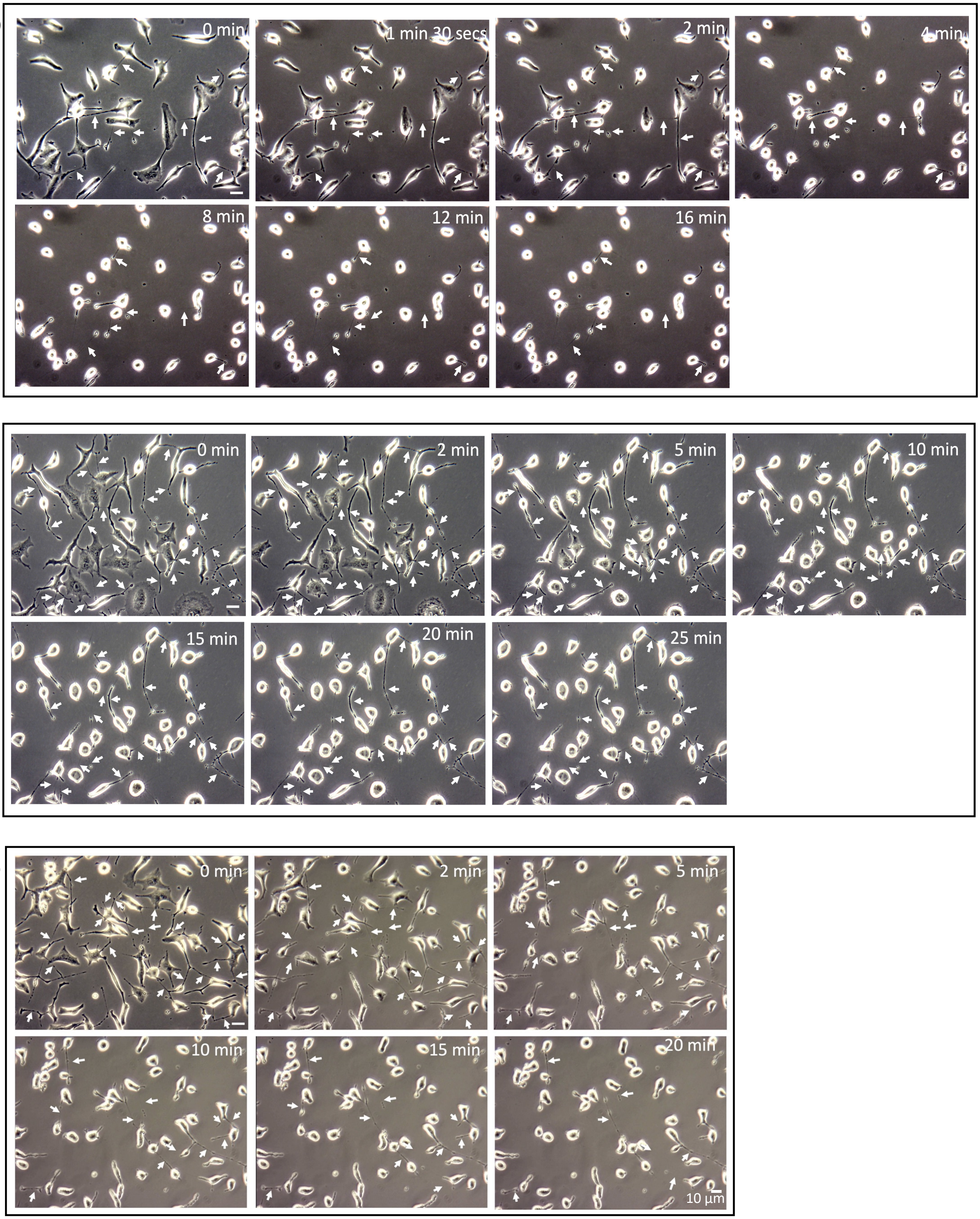


**Supplementary figure 1A. EGF, HGF and EGF+HGF-induced TNTs are retained post-trypsinisation** Representative phase contrast timelapse image sequences demonstrating (i) EGF (ii) HGF and (iii) EGF+HGF-induced TNTs are retained at least 16, 25 and 20 minutes post-trypsinisation respectively. Timelapse movies were recorded at 10x magnification. (n=3)

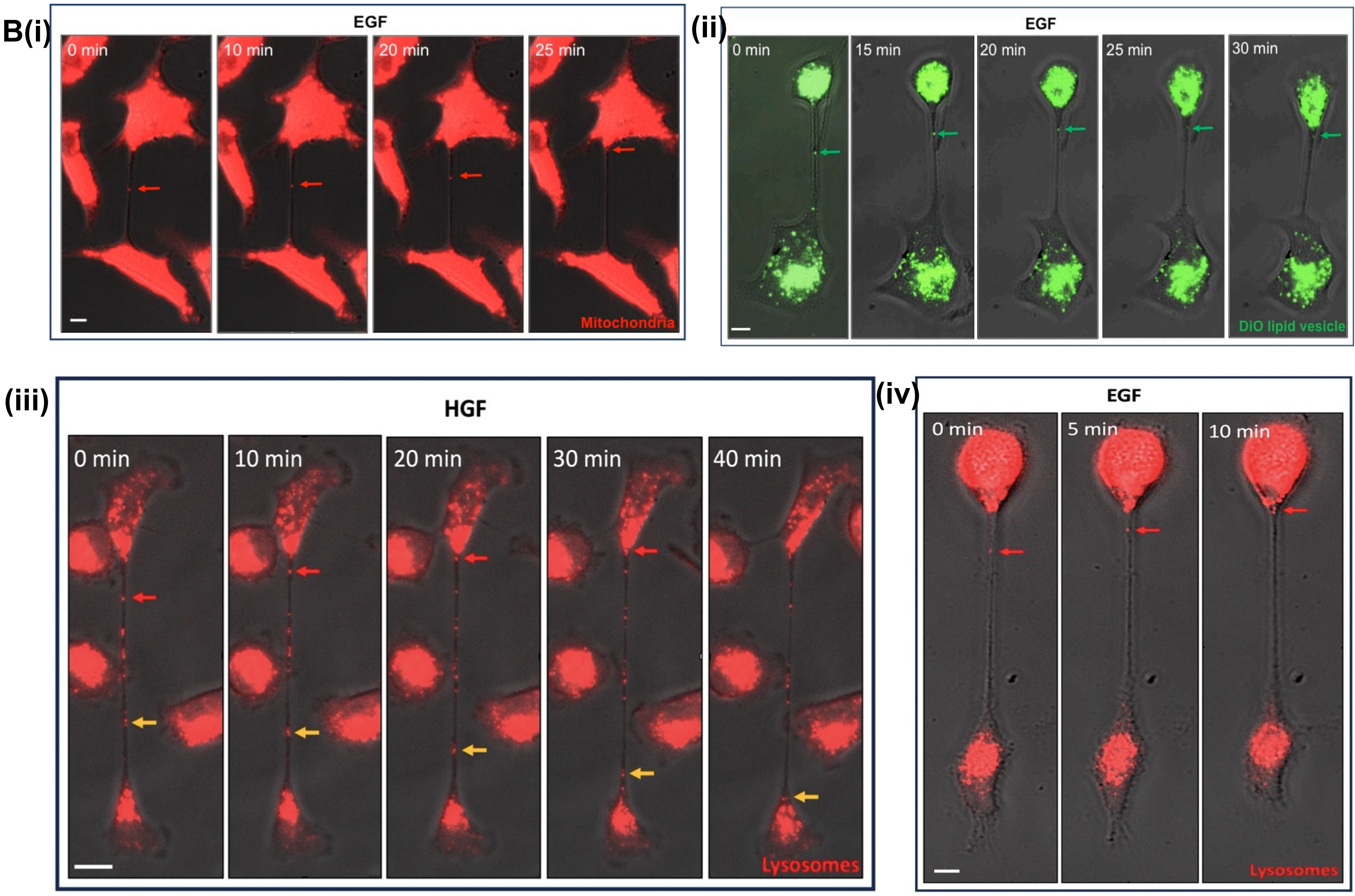


**Supplementary figure 1B. TNTs facilitate intercellular trafficking of mitochondria, DiO-labelled lipid vesicles and lysosomes in A549 cells (i)** Representative merged phase contrast and fluorescent images of (i) mitochondria (red arrow) being transported within an EGF-induced TNT. **(ii)** a DiO-labelled vesicle (green arrow) being trafficked within an EGF-induced TNT **(iii)** lysosomes (red and yellow arrows) being bidirectionally transported within a HGF-induced TNT. **(iv)** lysosomes (red arrow) being bidirectionally transported within a HGF-induced TNT. Timelapse movies were obtained at 20x magnification. Scale bar: 5 µm. (n=3)

A (i)

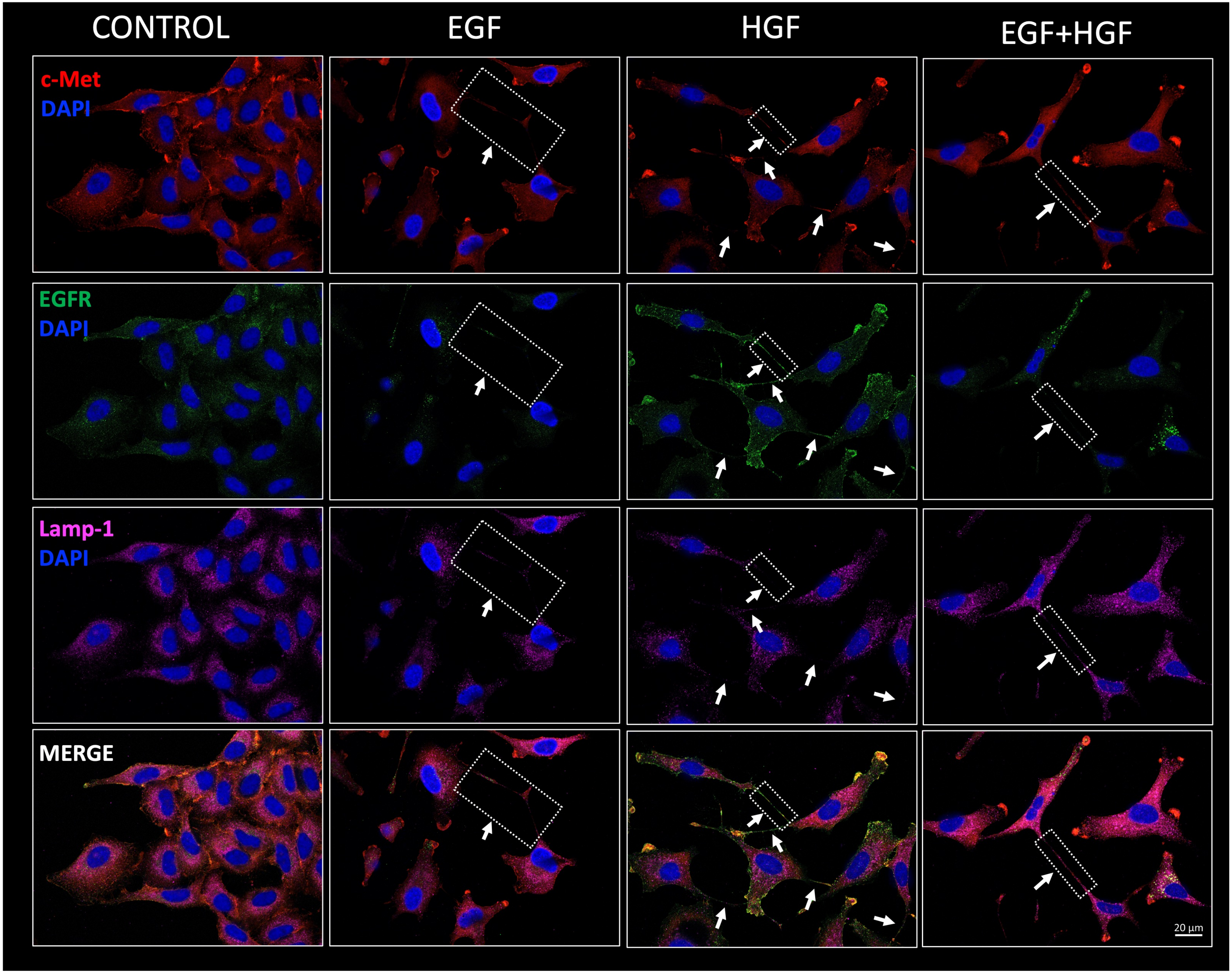
CONTROL
EGF
HGF
EGF+HGF
c-Met
DAPI
EGFR
DAPI
Lamp-1
DAPI
MERGE
20 µm

(ii)

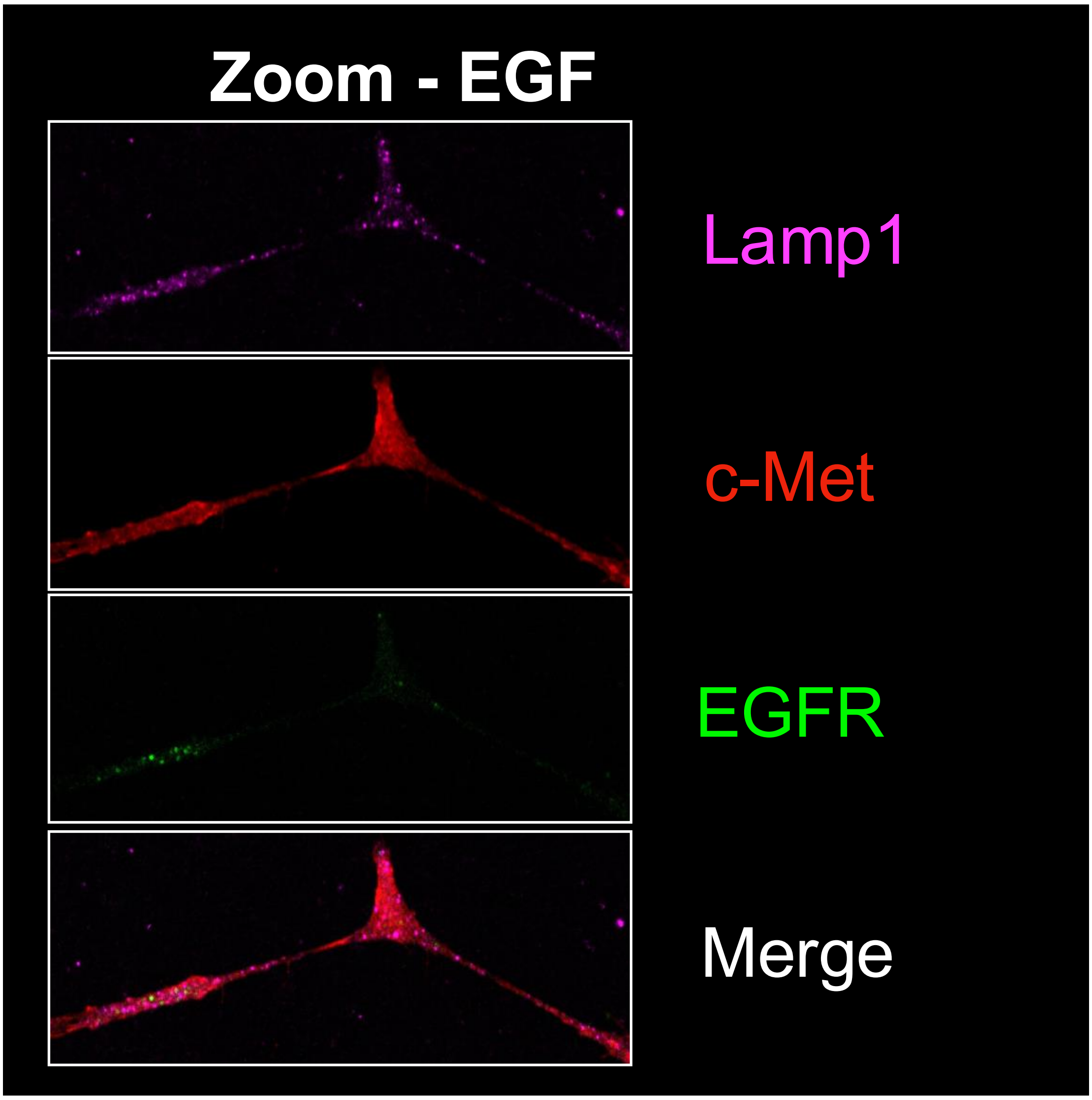
Zoom - EGF
Lamp1
c-Met
EGFR
Merge

(iii)

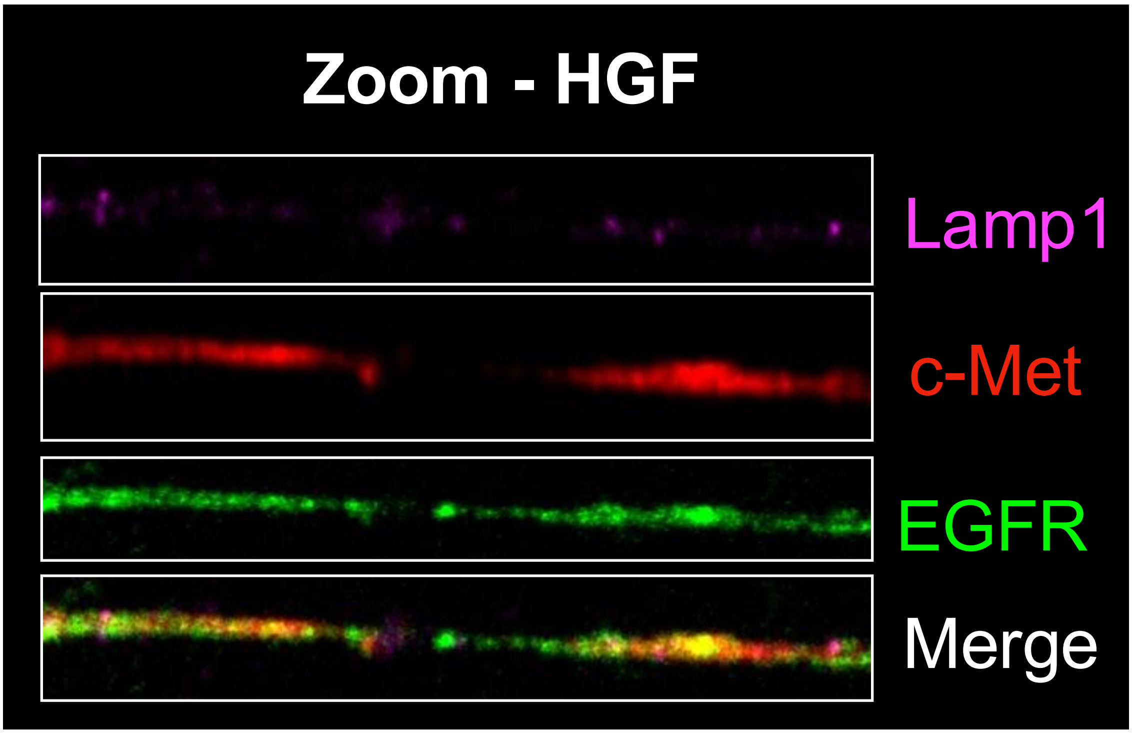
Zoom - HGF
Lamp1
c-Met
EGFR
Merge

(iv)

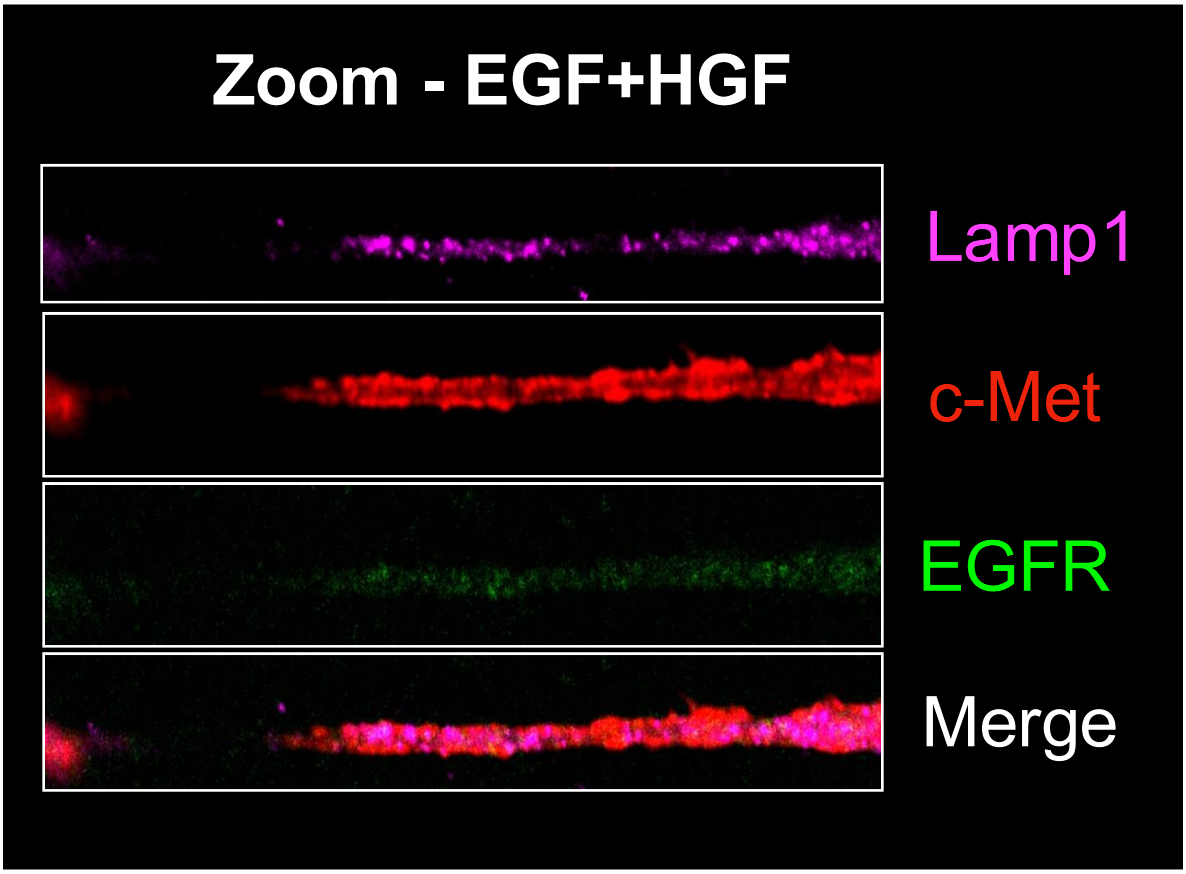


**Supplementary figure 2A. Lamp-1 localised to EGF, HGF and EGF+HGF-induced TNTs in A549 cells, however it did not co-localise with the EGFR and c-Met receptor.** (i) Representative confocal images illustrating the localisation of Lamp-1 (magenta), c-Met (red) and EGFR (green) in unstimulated cells, or cells stimulated with EGF,HGF, and EGF+HGF. Lamp-1 did not co-localise with either EGFR or c-Met within TNTs, with the receptors and Lamp-1 puncta remaining discrete along the length of the TNTs. White arrows indicate TNT structures. Zoom panels show magnified views of the dotted regions of (ii) EGF (iii) HGF and (iv) EGF+HGF-induced TNTs expressing (top to bottom) Lamp-1 (magenta), c-Met (red), EGFR (green) and merged. Images obtained at x40 magnification using a confocal microscope. (n=3)

B (i)

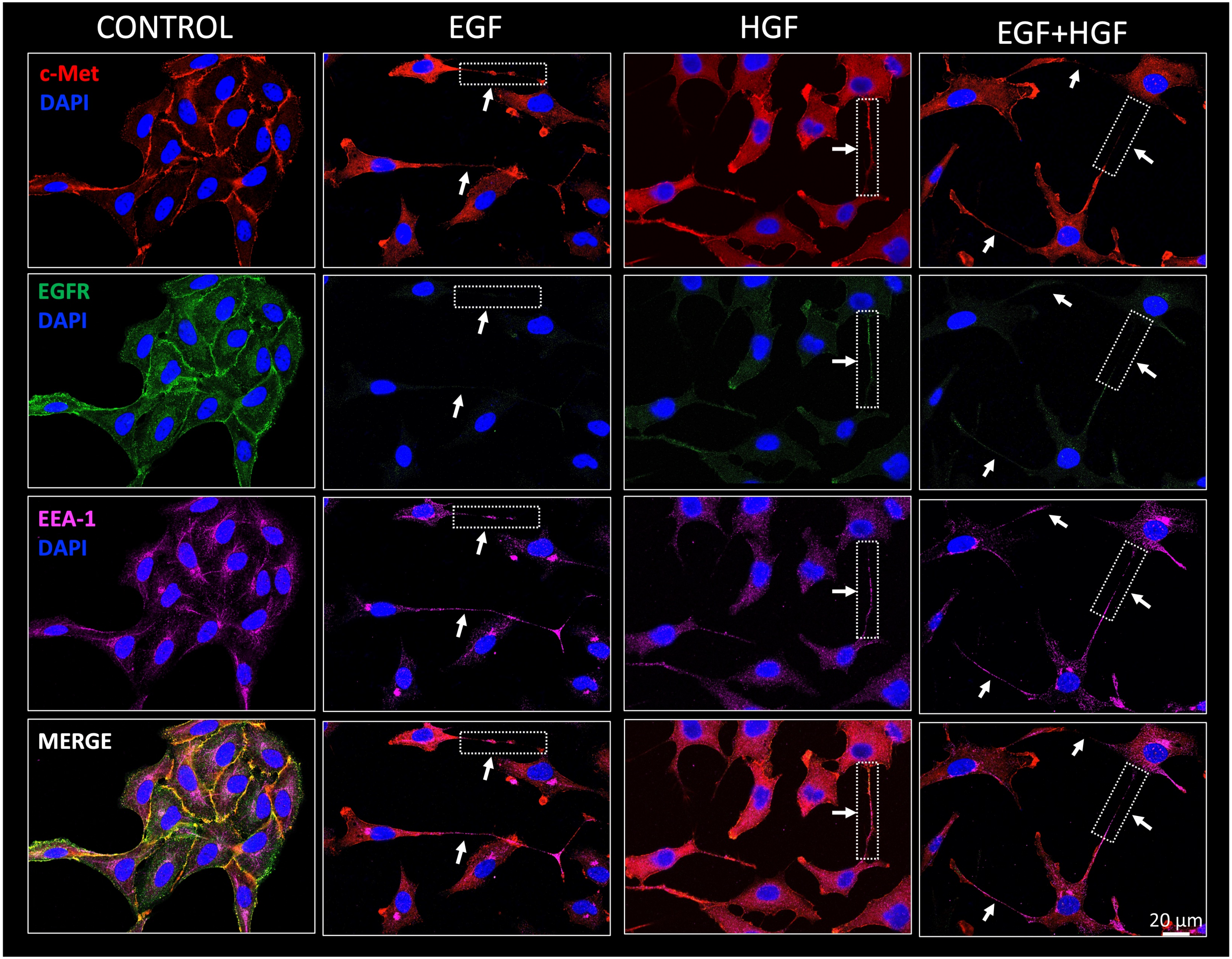
CONTROL
EGF
HGF
EGF+HGF
c-Met
DAPI
EGFR
DAPI
EEA-1
DAPI
MERGE
20 µm

(ii)

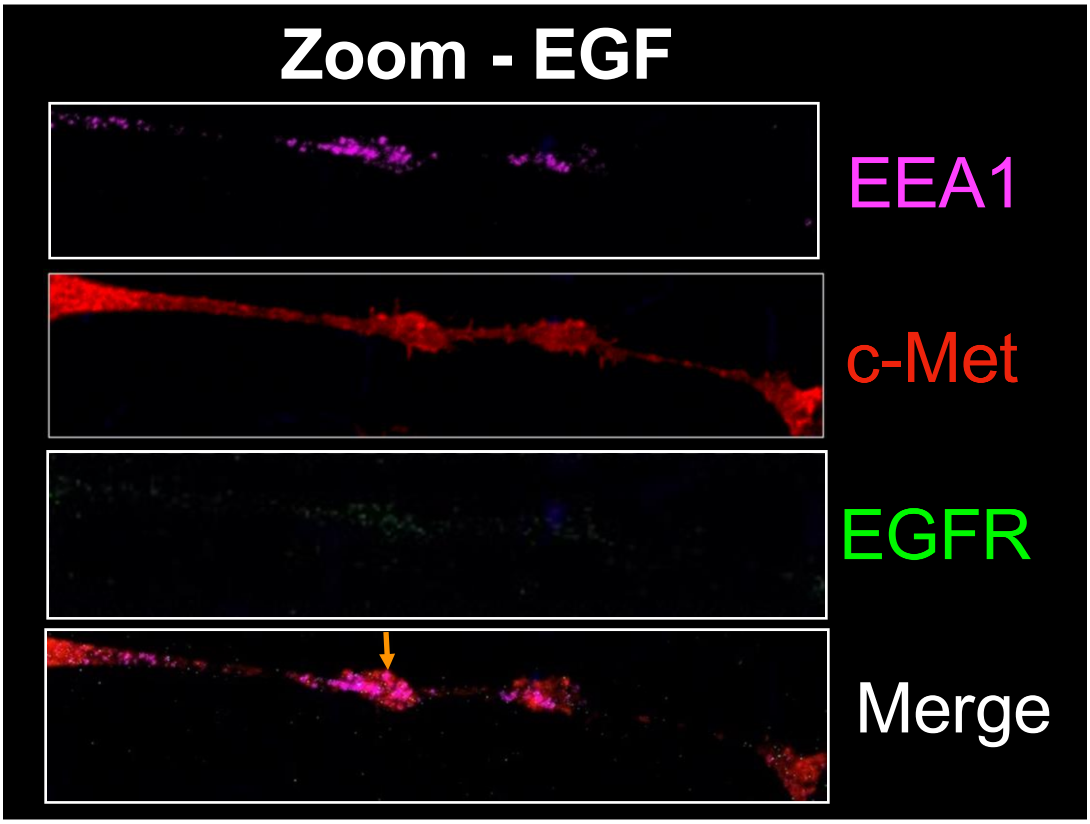


(iii)

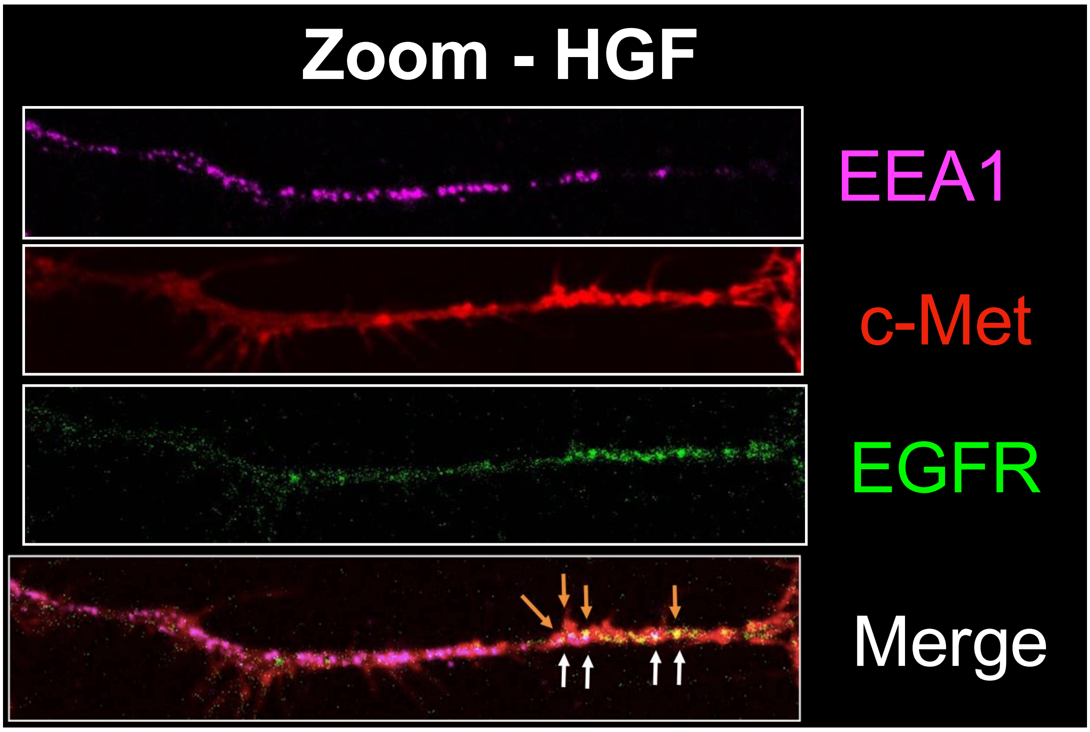

(iv)

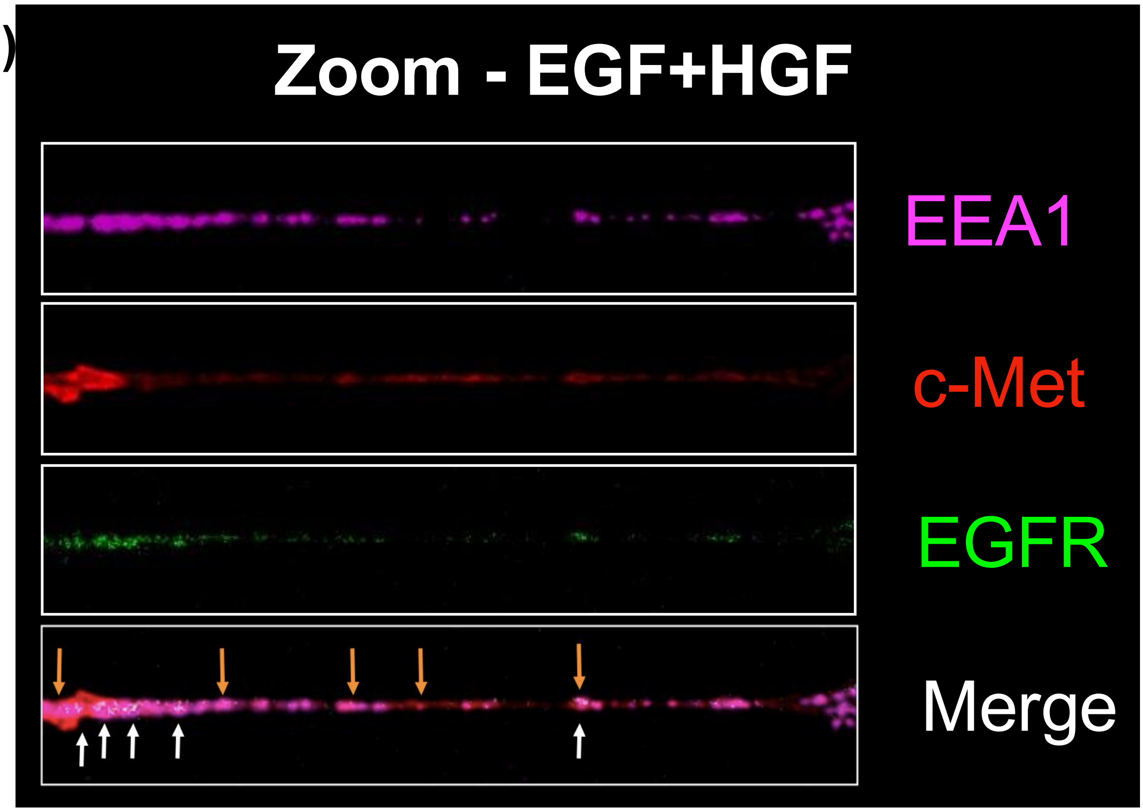


**Supplementary figure 2B. EEA1 is a novel component of HGF, EGF and EGF+HGF-induced TNTs in A549 cells and occasionally co-localised with the EGFR and c-Met receptor.** (i) Representative confocal images showing the localisation of EEA-1 (magenta), c-Met (red) and EGFR (green) in unstimulated cells, or cells stimulated with HGF, EGF, and EGF+HGF. In TNTs induced by EGF, HGF, and EGF+HGF, EEA1 was detected along the TNTs, where it formed punctate structures. Co-localisation between EEA1 and EGFR/c-Met was observed in some TNTs, but this was not consistent across all instances. White arrows indicate TNT structures. Zoom panels show magnified views of the dotted regions of (ii) EGF (iii) HGF and (iv) EGF+HGF-induced TNTs expressing (top to bottom) EEA1 (magenta), c-Met (red), EGFR (green) and merged. Co-localisation between EEA1/c-Met is indicated using orange arrows and co-localisation between EEA1/EGFR is indicated using white arrows. Images obtained at x40 magnification using a confocal microscope. (n=3)
